# Diethylcarbamazine elicits Ca^2+^ signals through TRP-2 channels that are potentiated by emodepside in *Brugia malayi* muscles

**DOI:** 10.1101/2023.04.10.536248

**Authors:** Paul D. E. Williams, Sudhanva S. Kashyap, Alan P. Robertson, Richard J. Martin

## Abstract

Filarial nematode infections are a major health concern in several countries. Lymphatic filariasis is caused by *Wucheria bancrofti* and *Brugia spp.* affecting over 120 million people. Heavy infections can lead to elephantiasis having serious effects on individuals’ lives. Although current anthelmintics are effective at killing the microfilariae in the bloodstream, they have little to no effect against adult parasites found in the lymphatic system. The anthelmintic diethylcarbamazine is one of the central pillars of lymphatic filariasis control. Recent studies have reported that diethylcarbamazine can open Transient Receptor Potential (TRP) channels on the muscles of adult female *Brugia malayi* leading to contraction and paralysis. Diethylcarbamazine has synergistic effects in combination with emodepside on *Brugia* inhibiting motility: emodepside is an anthelmintic that has effects on filarial nematodes and is under trials for treatment of river blindness. Here we have studied the effects of diethylcarbamazine on single *Brugia* muscle cells by measuring the change in Ca^2+^ fluorescence in the muscle using Ca^2+^-imaging techniques. Diethylcarbamazine interacts with the TRPC orthologue receptor TRP-2 to promote Ca^2+^ entry into the *Brugia* muscle cells which can activate SLO-1 Ca^2+^ activated K^+^ channels, the putative target of emodepside. A combination of diethylcarbamazine and emodepside leads to a bigger Ca^2+^ signal than when either compound is applied alone. Our study shows that diethylcarbamazine targets TRP channels to promote Ca^2+^ entry that is increased by emodepside activation of SLO-1 channels.

## Introduction

Lymphatic filariasis (elephantiasis) is a neglected tropical disease caused by filarial nematode parasites that affects more than 120 million people worldwide, with 40 million people suffering from disfiguration or disability [1]. Lymphatic filariasis is caused by the adults of the filarial parasites *Wuchereria bancrofti, Brugia malayi,* and *Brugia timori*. These parasites are transmitted between hosts through biting insects, and the adults live within the lymphatic vessels. In severe cases, the parasites block lymphatic ducts resulting in swelling of limbs and coarsening of the skin resulting in the condition known as elephantiasis. The swelling of limbs can result in disabilities, societal rejection and prevent individuals from working. There are no effective vaccines against filarial parasites, nor have the measures to control the spread by their vectors been adequate.

Control of lymphatic filariasis relies on the use of chemotherapeutics to target and kill the microfilaria to prevent transmission between individuals by biting insects. The anthelmintics used to treat lymphatic filariasis include the: benzimidazoles (albendazole) that bind to β-tubulin, inhibiting microtubule formation as well as metabolism [2]; macrocyclic lactones (ivermectin), which target the glutamate-gated chloride channels [3] and; diethylcarbamazine (diethylcarbamazine), a compound whose mode of action is not well characterized but have been reported to interact with Transient Receptor Potential (TRP) channels [4, 5]. Each of these anthelmintics may be administered alone or in combination in Mass Drug Administration (MDA) programs [6, 7]. However, use of ivermectin or diethylcarbamazine in individuals who suffer from loiasis, or onchocerciasis is not recommended by the CDC or WHO because of complications that could lead to aggravation of *Onchocerca* induced eye disease or loiasis encephalopathy and death. Although effective at killing and clearing microfilaria, none of these anthelmintics kill all of the adult worms: the surviving parasites live for 6-8 years producing microfilaria.

Historically, diethylcarbamazine had been understood to act on the host immune system rather than the parasite itself [8-10]. However, recent studies have shown that diethylcarbamazine can stimulate nematode TRP channels, including the TRPC orthologue TRP-2 and the TRPM orthologues GON-2 and CED-11 [4, 5]. In *Brugia* muscle cells, the effect of diethylcarbamazine opening TRP channels is an inward depolarizing current and contraction with spastic paralysis. The inward current produced by diethylcarbamazine is followed by an outward current produced by activation of Ca^2+^ activated SLO-1 K^+^ channels. The anthelmintic emodepside, which activates nematode SLO-1 channels [11-14] is a broad spectrum antiparasitic drug that has been reported to target and kill adult filaria. However, the *in vivo* potency of emodepside against some filarial parasites, like *Brugia,* is limited. Combinations of diethylcarbamazine and emodepside may promote increased efficacy: previous studies have highlighted the synergism between diethylcarbamazine and emodepside on muscle membrane potentials in the soil transmitted helminth *Ascaris suum* [12]; and the long-term muscle paralysis in *Brugia malayi* which is dependent on TRP-2 channels [13, 15].

In this study we have utilized Ca^2+^ imaging to identify the role of Ca^2+^ in mediating diethylcarbamazine signaling and the role of the TRP-2 in promoting Ca^2+^ entry. We show that TRP-2 is a major source of diethylcarbamazine stimulated Ca^2+^ entry as inhibition of the channels with SKF96365 and dsRNA knockdown inhibits the Ca^2+^ signal. We have tested the effects of arachidonic acid (AA) and miconazole and have observed that they increase Ca^2+^ entry into the cytoplasm and TRP channel activation. Finally, we identify a synergistic relationship between diethylcarbamazine and emodepside in potentiating the Ca^2+^ signal that explains the enhanced paralysis observed in motility assays by disrupting the homeostasis of Ca^2+^ in the muscle cell. These results support previous observations that diethylcarbamazine interacts with TRP channels and that the combination of diethylcarbamazine and emodepside has a potential for the treatment of individuals infected with *Brugia malayi*.

## Materials and Methods

### Brugia supply and maintenance

Only female *Brugia malayi* worms were used for the study. Live adult female *B. malayi* were shipped overnight from the NIH/NIAID Filariasis Research Reagent Resource Centre (FR3; College of Veterinary Medicine, University of Georgia, Athens, USA). *B. malayi* were maintained in non-phenol red Hyclone Roswell Park Memorial Institute (RPMI) 1640 media (Cytiva, USA) containing 10% heat-inactivated FBS (Fisher-Scientific) and 1% penicillin-streptomycin (Life Technologies, USA). Parasites were separated individually into a 24 well microtiter plate containing 2mL of the RPMI media. Parasites were held in an incubator set to 37°C and 5% CO_2_. All parasites were used up to 5 days post-delivery.

### Dissection of B. malayi

Dissection of *B. malayi* was performed as previously described [16-18]. All dissections were performed at room temperature. Briefly, worms were cut into 1 cm pieces from the anterior region and single sections placed into the recording chamber filled with *B. malayi* bath solution (23 mM NaCl, 110 mM NaAc, 5 mM KCl, 1 mM CaCl_2_, 4 mM MgCl_2_, 5 mM HEPES, 11 mM D-glucose, 10 mM Sucrose, pH 7.2 using NaOH, ∼320 mOsmol). The base of the chamber was a coverslip (24 x 50 mm) coated with a thin layer of Sylgard. The body piece was immobilized gluing each end to the Sylgard pad using Glushield cyanoacrylate glue (Glustitch, BC, Canada) and immobilized by creating a wall down one side under the dissecting microscope. The body piece was cut longitudinally using a tungsten needle, and the ‘muscle flap’ was glued to the coverslip along the cut edge exposing the muscle cells. The intestines and the uterus were removed using fine forceps and the prep was washed with bath solution to remove any eggs or debris.

### Fluo-4 injection

To record Ca^2+^ signals, muscles were injected with 5 µM Fluo-4 penta-potassium salt (ThermoFisher Scientific, USA) as previously described and viewed with DIC optics on a Nikon Eclipse TE300 inverted light microscope (400×) [19]. Briefly, the dissected worms were treated with 2mg/mL collagenase (Type 1A, Gibco) 20-30 seconds and washed several times with buffer to remove excess collagenase. Patch pipettes were pulled from capillary glass and fire polished. The pipettes were filled with pipette solution (120 mM KCL, 20 mM KOH, 4 mM MgCl_2_, 5 mM TRIS, 0.25 mM CaCl_2_, 4 mM NaATP, 5 mM EGTA and 36 mM sucrose (pH 7.2 with KOH ∼ 315-330 mOmol). Fluo-4 was added to the pipette solution at the start of each recording day at a concentration of 5 µM and was kept in a dark environment to prevent degradation of the dye. Pipettes with a resistance of 1.5-3MΩ were used. Giga ohm seals were formed before breaking the membrane by suction. After breaking in, cells were left to allow the Fluo-4 solution to diffuse the entire muscle cell Fig.1A (∼5-10 minutes).

**Figure 1:**
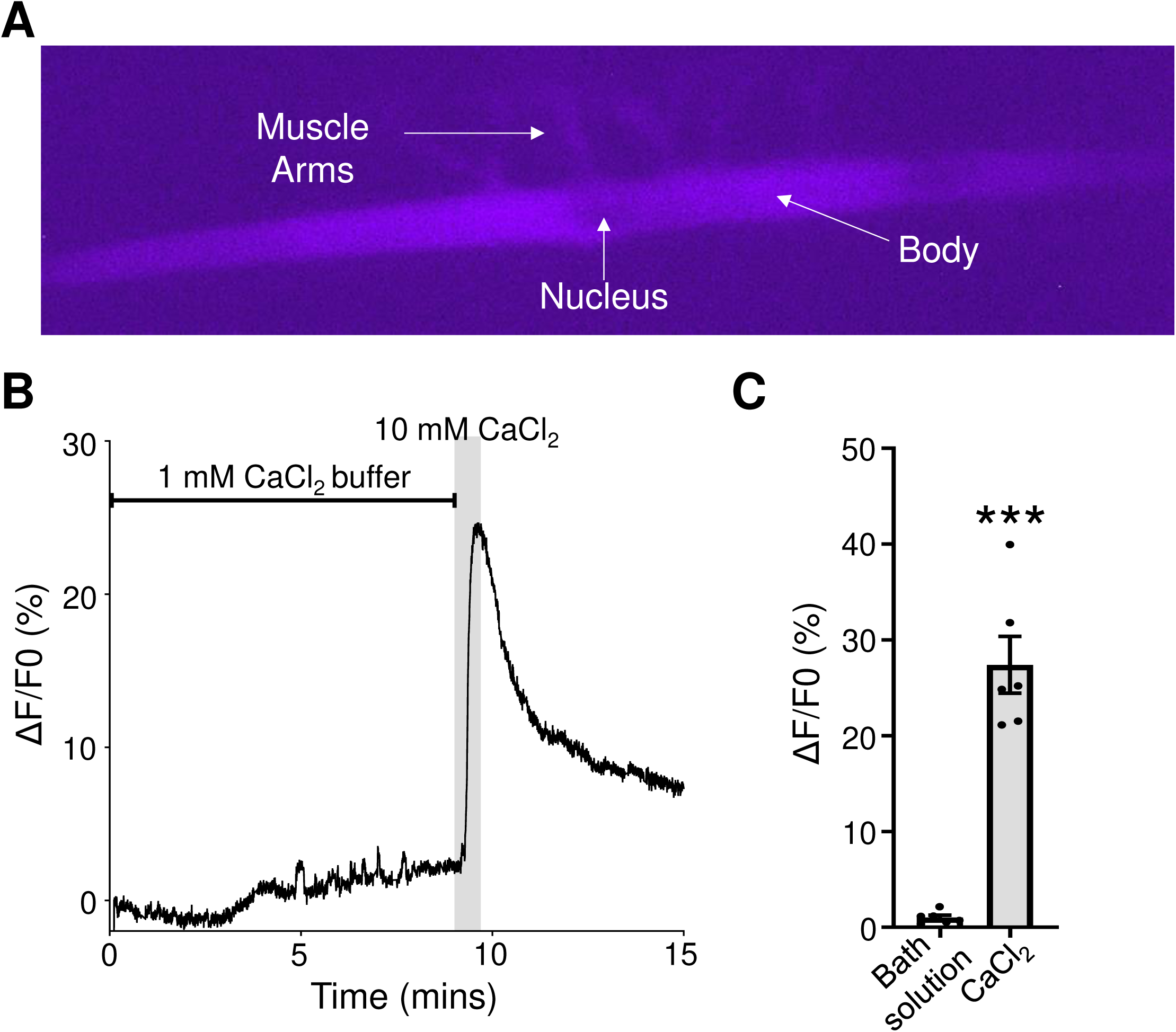
Fluo-4 induced Ca^2+^ signals in *Brugia* muscles. A) Micrograph of a fluorescing dissected *Brugia* muscle injected with 5 µM Fluo-4 under blue light. Key structures of the body, nucleus and muscle arms are indicated with the arrows. B) Representative control trace of Fluo-4 fluorescence in a muscle being washed with bath solution containing 1 mM CaCl_2_ followed by an increase to 10 mM CaCl_2_ for 1 minute (grey box). C) Total amplitudes of Fluo-4 fluorescence in response to constant 1mM CaCl_2_ bath solution perfusion (black bar) and to 10 mM CaCl_2_ (grey bar). *** Significantly different to 1 mM CaCl_2_ bath solution (Bath solution vs CaCl_2_ *P <* 0.001 *t* = 8.817, *df* = 5, paired *t*-test. *n* = 6 muscles. All values represented as means ± SEM.

### Ca^2+^ imaging

All recordings were performed with a Nikon Eclipse TE300 microscope (20X/0.45 Nikon PlanFluor objective), fitted with a Photometrics Retiga R1 Camera (Photometrics, Surrey, BC, Canada). Light control was achieved using a Lambda 10-2 two filter wheel system with a shutter controller (Lambda Instruments, Switzerland). Filter wheel one was set on a green filter between the microscope and camera. Filter wheel two set on the blue filter between a Lambda LS Xenon bulb light box which delivered light via a fiber optic cable to the microscope (Lambda Instruments, Switzerland). Blue light emission was controlled using a shutter. All Ca^2+^ signal recordings were acquired and analyzed using MetaFluor 7.10.2 (MDS Analytical Technologies, Sunnyvale, CA), Exposure times were 150ms with 2x binning. Maximal Ca^2+^ signal amplitudes (ΔF %) for all stimuli applied were calculated using the equation F1-F0/F0 x 100 where F1 is the fluorescent value and F0 is the baseline fluorescent value, which was determined as the value immediately before the stimulus was applied to the sample for all recordings analyzed. Representative traces were generated using the same formula, with F0 being the value before the application of the first stimulus. For 1 mM CaCl_2_ bath solution control experiments (Fig. 1B & C) F0 was determined to be the first value of the recording. Rise times were calculated by normalizing the trace during stimulus exposure until the application of the 10 mM CaCl_2_ positive control, with the lowest fluorescence value being represented by 0% and the highest being 100%. The peak time was calculated by subtracting the time when the stimulus was applied from the time the signal reached 100%.

### Application of compounds

The preparation was constantly perfused with all solutions being delivered to the chamber under gravity feed through solenoid valves controlled with a VC-6 six channel Valve Controller (Warner Instruments, Hamden, CT) through an inline heater set at 37°C (Warner Instruments, Hamden, CT) at a rate of 1.5mL/min. New preparations were perfused with bath solution containing 1 mM CaCl_2_ before being exposed to either 30 µM diethylcarbamazine (DEC) for 5 minutes, 10 µM arachidonic acid (AA) for 5 minutes, 10 µM miconazole (MIC) for 5 minutes or 1 µM emodepside (Emo) for 5 minutes. Exposure to diethylcarbamazine and SKF96365 was achieved by exposing muscle cells to 10 µM SKF96365 alone (SKF) for 5 minutes, followed by 10 µM SKF96365 and 30 µM diethylcarbamazine for 5 minutes (SKF + DEC), and finally to 30 µM diethylcarbamazine (DEC) alone for 5 minutes giving a total exposure time of 15 minutes. For diethylcarbamazine and emodepside combination experiments, muscle cells were exposed to 30 µM diethylcarbamazine (DEC) for 5 minutes and then 30 µM diethylcarbamazine + 1 µM emodepside (DEC + Emo) for 5 minutes, for a total of 10 minutes. To ensure that each of the preparations were viable after being exposed to any of the stimuli tested, all samples were subjected to 10 mM CaCl_2_ for 1 minute to act as a positive control. Muscles that elicited Ca^2+^ signals to 10 mM CaCl_2_ (> 10 %) were considered viable while those that failed to elicit response (< 10 %) were classified as non-viable.

### Synthesis and delivery of dsRNA

*Trp-2* target T7 promoter labeled primers were amplified using the primers *trp-2f2, trp-2f2t7, trp-2r2* and *trp-2r2t7* previously described (Supplementary Table 1) [4, 15]. Non-specific T7 labelled *LacZ* dsRNA constructs were amplified using the primers *LacZf, LacZr, LacZft7*, and *LacZrt7* (Supplementary Table 1) [15]. Amplification was performed from sequence verified cDNA templates using a Techne^TM^ PRIMEG cycler (Bibby Scientific Limited, UK) with the following cycling conditions: 95°C x 5 min, 35 x (95°C x 30 s, 55 °C x 30 s, 72 °C x 1 min), 72 °C x 10 min. dsRNA probes were synthesized [18, 20] using the T7 RiboMAX^TM^ Express RNAi kit (Promega, USA) according to manufacturer’s instructions (Supplementary Table 2). The concentration of dsRNA was assessed using a spectrophotometer. Adult female *B. malayi* were soaked in 30 µg/mL of LacZ or *trp-2* dsRNA for 4 days before being used for Ca^2+^ imaging experiments.

### Analysis of transcript levels

cDNA from dsRNA-treated worms were amplified using the following target *trp-2* and reference gene (*Bma*-*gapdh*) primers: *trp-2f2, trp-2r2*, SSK5F, and SSK5R (Supplementary Table 1) [4, 15]. These genes were amplified in triplicate by quantitative real-time PCR (qPCR) using the CFX96 TouchTM Real-Time PCR Detection System and SsoAdvancedTM Universal SYBR® Green Supermix (Bio-Rad, USA). The cycling conditions used were 95 °C × 10 min, 40 × (95 °C × 10 s, 55 °C × 30 s). PCR efficiencies were calculated using the CFX96 Software Suite (Bio-Rad, USA). The relative quantification of target gene knockdown was estimated by the ΔΔCt method [21].

### Chemicals

Emodepside was procured from Advanced ChemBlocks, SKF96365 was procured from Tocris; Sigma Aldrich supplied all other chemicals. All compounds were dissolved in either water or DMSO and diluted in bath solution to obtain final concentrations. DMSO final concentration, 0.01%

### Statistical analysis

Statistical analysis of all data was done using GraphPad Prism 9.0 (Graphpad Software, Inc., La Jolla, CA, USA). To ensure reproducibility, we repeated our experiments: the numbers of samples, the concentrations, and durations of applications of either diethylcarbamazine, arachidonic acid, miconazole, SKF96365 and emodepside are provided in the methods and legends of the figures. Analysis of Ca^2+^ amplitudes were done using either unpaired or paired student *t*-tests with *P* values <0.05 considered as significant using Prism Graphpad version 9.0 software.

## Results

### Diethylcarbamazine stimulates a detectable Ca^2+^ response in Brugia muscles

Ca^2+^ signals can be recorded in the muscles of *Brugia malayi* by directly injecting a fluorescent dye into the cell via a patch pipette [19]. Here we use Fluo-4, an analogue of Fluo-3, which has increased fluorescence excitation and higher fluorescence signal levels according to the manufacturer. We loaded individual muscle cells with Fluo-4 until we were able to see key structures of the muscle cell including the muscle arms, nucleus and the cell body (Fig. 1A). We exposed muscle cells to bath solutions containing 1 mM CaCl_2_ and observed small (1.0 %, ± 0.3 %, *n* = 6) fluctuations in the fluorescent signal (Fig. 1B and C (black bar)) due miniature end-plate potentials that are seen in the *Brugia* muscle cells. To verify that our muscles are physiologically active, we increased the CaCl_2_ concentration by applying 10 mM CaCl_2_ to the preparation. We observed rapid large reversible increases in fluorescence (27.4 % ± 3.0 %, *n* = 6) when 10 mM CaCl_2_ was applied, and that fell when it was removed (Fig 1B and C; grey bar).

We have reported previously that diethylcarbamazine inhibits motility and produces an initial spastic paralysis in *B. malayi* that is associated with inward currents on the muscle [4]. This inward current was shown to be dependent on cation permeable Transient Receptor Potential (TRP) channels [4]. We hypothesized that the activation of these channels would allow the entry of Ca^2+^ into the cell, which we could detect using our Ca^2+^ imaging protocol. We exposed muscles cells loaded with Fluo-4 to 30 µM diethylcarbamazine for 5 minutes and observed a characteristic increase in the Ca^2+^ signal which decreased when the diethylcarbamazine was removed (Fig. 2A). The average increase in fluorescence to diethylcarbamazine was 15% (15.4 %, ± 4.2 %, *n* = 7: Fig. 2B; brown bar). The cells were then challenged with 10 mM CaCl_2_ to ensure they were functioning physiologically, and we observed an increase in fluorescence (Fig. 2A & B; grey bar, 41.4%, ± 11.9 %, *n* = 7). We see that diethylcarbamazine simulates a reversible rise in sarcoplasmic Ca^2+^ in *Brugia* muscle cells that is consistent with the opening of TRP channels that are permeable to Ca^2+^.

**Figure 2:**
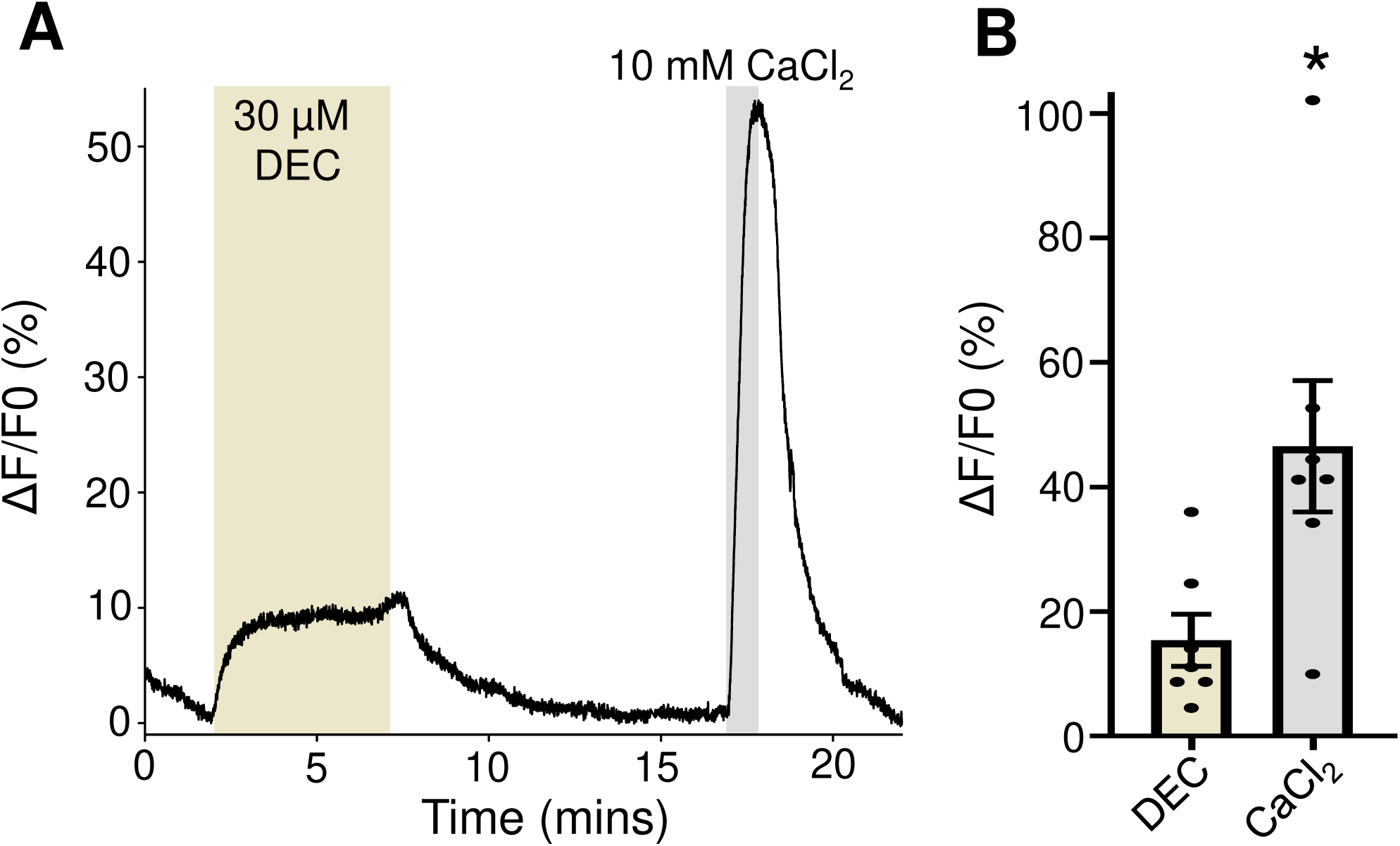
Diethylcarbamazine stimulates increases in Ca^2+^ in adult *Brugia* muscles. A) Representative Ca^2+^ trace in response to 30 µM diethylcarbamazine (brown box) for 5 minutes followed by 10 mM CaCl_2_ control for 1 minute (grey box). B) Total amplitudes of Fluo-4 fluorescence to 30 µM diethylcarbamazine (DEC) (light brown bar) and 10 mM CaCl_2_ (grey bar). * Significantly different to CaCl_2_ (diethylcarbamazine (DEC) vs CaCl_2_) *P <* 0.017, *t* = 3.259, *df* = 6, paired *t*-test. *n* = 7 muscles. All values represented as means ± SEM.

### Arachidonic acid and miconazole that stimulate TRP channels also promote Ca^2+^ entry

Nematodes produce poly unsaturated fatty acids (PUFAs) by converting arachidonic acid into biologically active and inactive PUFAs via ω-hydroxylases and epoxygenases enzymes. Verma *et al.,* [4] showed that arachidonic acid has a similar effect on the amplitude of the inward current on *B. malayi* muscles as diethylcarbamazine, but the signal was slower than diethylcarbamazine in onset and reaching a peak suggesting that arachidonic acid metabolites were responsible for the activation of the TRP channels. Here we investigated the role of arachidonic acid on the Ca^2+^ signal by exposing the muscles to arachidonic acid. 10 µM arachidonic acid stimulated a Ca^2+^ signal that had a similar profile to that of diethylcarbamazine (Fig 3A; compare blue trace to black trace) and a similar overall amplitude (13.3 %, ±1.9 %, *n* = 6 vs: Fig 3B; blue bar). Although similar in amplitude, the arachidonic acid signals had a significantly slower rise to the peak of the Ca^2+^ signal compared to diethylcarbamazine (5.6 minutes ± 1.0 *n* = 6 vs 2.6 minutes, ± 0.8, *n* = 7; Fig 3C; blue bar), a phenotype that mimics the slower electrophysiology responses to arachidonic acid [4].

**Figure 3:**
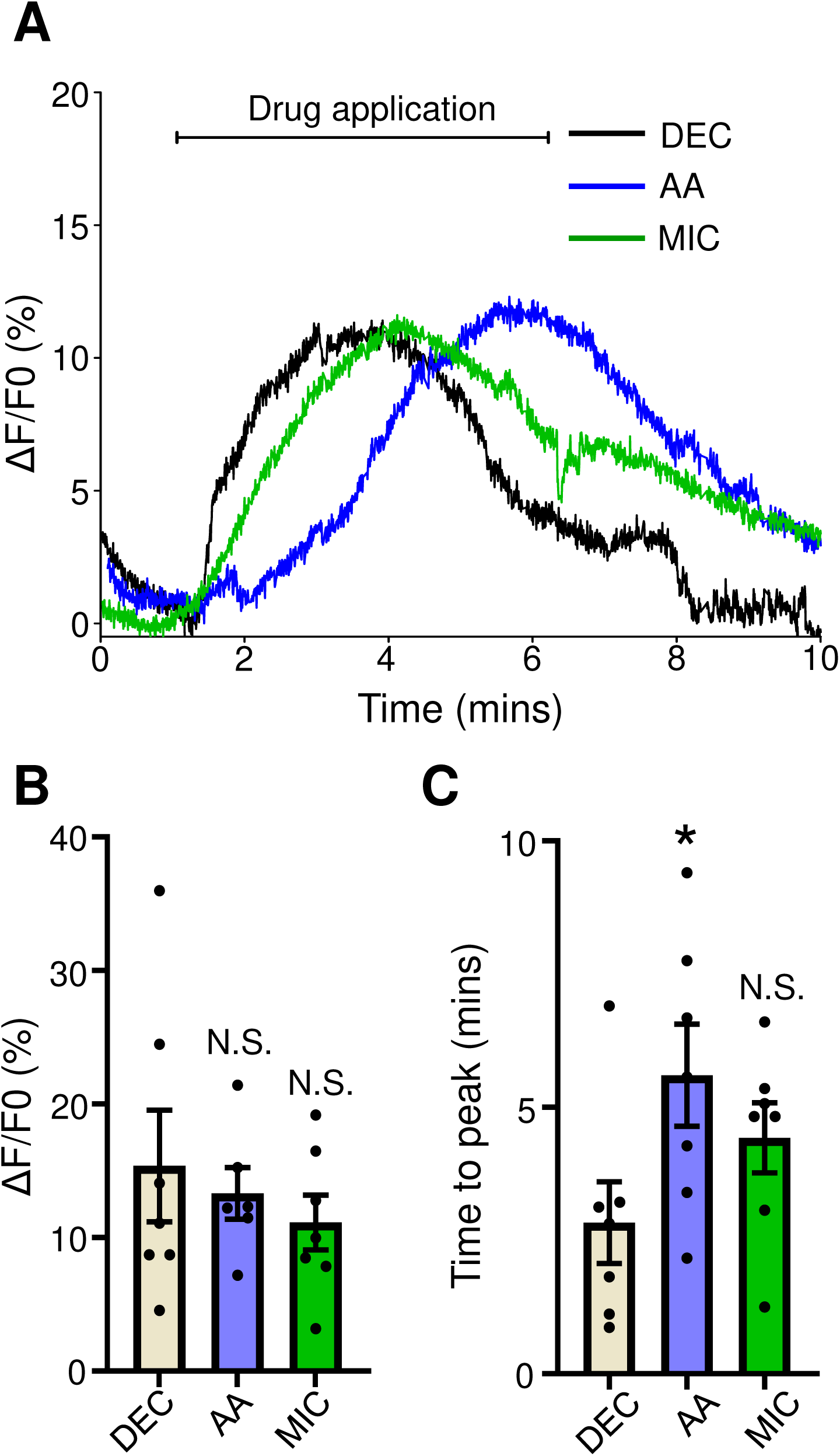
Arachidonic acid and miconazole stimulate Ca^2+^ signals. A) Representative individual traces of Ca^2+^ signals to either 30 µM diethylcarbamazine (black line), 10 µM arachidonic acid (blue line) and 10 µM miconazole (green line). The bar represents application of the compounds over 5 minutes. B) Total amplitudes for diethylcarbamazine (DEC; light brown bar), arachidonic acid (AA; blue bar) and miconazole (MIC; green bar). N.S. not significantly different to diethylcarbamazine: diethylcarbamazine (DEC) vs arachidonic acid (AA), *P <* 0.681, *t* = 0.422, *df* = 11 unpaired *t*-test; diethylcarbamazine (DEC) vs miconazole (MIC)), *P <* 0.381, *t* = 0.909, *df* = 12, unpaired *t*-test. C) Average time to maximum peak (100% of signal) for diethylcarbamazine (DEC; light brown bar), arachidonic acid (AA; blue bar) and miconazole (MIC; green bar). * Significantly different to diethylcarbamazine: diethylcarbamazine (DEC) vs arachidonic acid (AA), *P <* 0.044, *t* = 2.256, *df* = 12, unpaired *t*-test. N.S. not significantly different to diethylcarbamazine: diethylcarbamazine (DEC) vs miconazole (MIC), *P <* 0.142, *t* = 1.573, *df* = 1, unpaired *t*-test. *n* = 7 for diethylcarbamazine responses, *n* = 7 for arachidonic acid and *n* = 7 miconazole treatments. All values represented as means ± SEM.

Miconazole is an inhibitor of epoxygenase CYP450 enzymes that mimics the effects of arachidonic acid, which is explained by diverting the metabolism of arachidonic acid to active poly-unsaturated fatty acids (PUFAs) that open the TRP channels in *Brugia* muscle; like arachidonic acid, miconazole produces a slow opening of TRP channels current during its application [4].

We applied 10 µM miconazole to *Brugia* muscles and recorded the Ca^2+^ signals and observed Ca^2+^ signals with similar peak amplitudes to our responses to diethylcarbamazine and arachidonic acid (11.1 %, ± 2.1 %, *n* = 7: Fig 3A, green trace & B, green bar). The times to peak for the miconazole response were slower than our diethylcarbamazine responses (4.4 minutes ± 0.7, *n* = 7: Fig. 3C, green bar) although the difference did not reach statistical significance. Nonetheless, these results, taken together, further strengthen the idea that TRP channels play a key role in allowing the entry of Ca^2+^ into the muscle cells and that endogenous PUFAs can open TRP channels that yield similar Ca^2+^ amplitudes as the diethylcarbamazine mediated responses.

### TRP-2 mediates the diethylcarbamazine induced Ca^2+^ signal

The TRPC nematode orthologue channel, TRP-2, has been identified as a key channel in mediating diethylcarbamazine effects, particularly effects on muscle membrane currents and motility [4, 15]. To test if the observed diethylcarbamazine Ca^2+^ signal is mediated by Ca^2+^ entering through TRP-2 channels, we treated dissected muscles with the TRPC specific antagonist SKF96365, a compound which we have previously used to inhibit TRP channel activity in both *Brugia* and the gastrointestinal parasite *Ascaris suum* [4, 5]. We exposed muscles to 10 µM SKF96365 for 5 minutes and observed no changes in the Ca^2+^ profile (Fig 4A). We then exposed the muscle to 30 µM diethylcarbamazine in the presence of SKF96365 for an additional 5 minutes and observed that the Ca^2+^ signal was inhibited (1.9 %, ± 0.8 %, *n* = 9: Fig 4A & B light brown bar). We also noticed that the inhibition was maintained after washing the preparation in the continued presence of 30 µM diethylcarbamazine for a final 5 minutes and that the effects of SKF96365 were not readily reversed (Fig. 4A). We observed no inhibition to the control CaCl_2_ signal (33.2 %, ± 7.0 %, *n* =9: Fig 4B grey bar). This suggests that the TRPC channel TRP-2 is a major source of the diethylcarbamazine stimulated Ca^2+^ signal in *Brugia* muscles.

**Figure 4:**
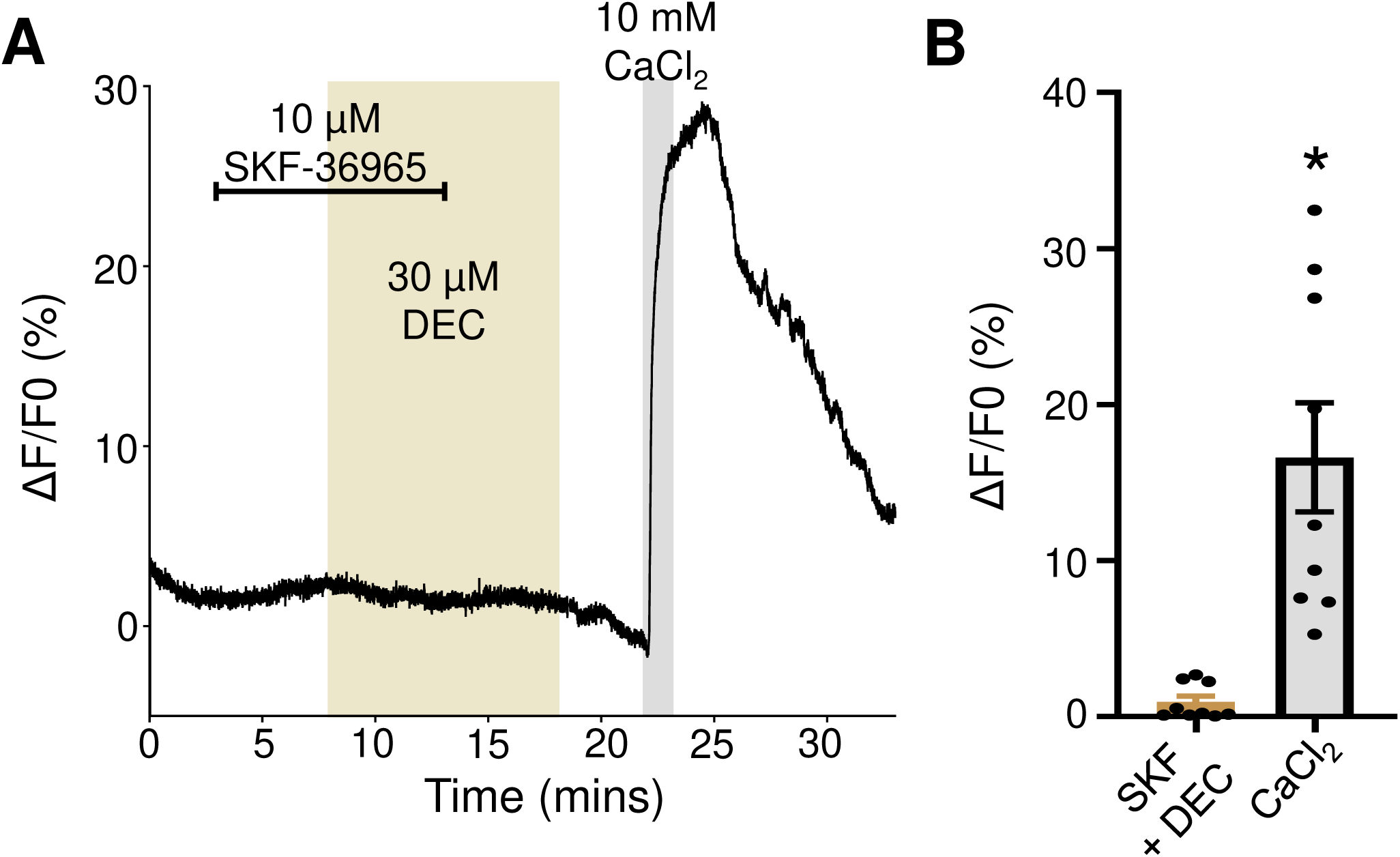
Inhibition of TRP-2 ablates diethylcarbamazine sensitivity. A) Representative trace showing application of 10 µM SKF96365 (light brown bar) for 5 minutes alone followed by combination of SKF96365 and of 30 µM diethylcarbamazine (light brown box) for 5 minutes. SKF96365 is removed leaving diethylcarbamazine alone for a final 5-minute exposure. 10 mM CaCl_2_ control was applied for 1 minute (grey box) as a positive control. B) Total Ca^2+^ amplitudes in response to SKF96365 + diethylcarbamazine (SKF + DEC) (light brown bar) and CaCl_2_ control (grey bar). * Significantly different to SKF96365 + diethylcarbamazine. SKF96365 + diethylcarbamazine (SKF + DEC) vs CaCl_2_*, P <* 0.002, *t* = 4.480, *df* = 8, paired *t*-test. *N* = 9 for diethylcarbamazine + SKF96365 responses and *n* = 9 for CaCl_2_ responses. All values represented as means ± SEM.

While SKF96365 is often used as a TRPC specific antagonist, it can affect other channels including the Ca^2+^ release-activated Ca^2+^ (CRAC) channel, STIM-1, voltage gated Ca_v_ channels and K^+^ channels [22-25]. To further test if the diethylcarbamazine mediated Ca^2+^ signal was due to TRP-2 activation, we used dsRNA to selectively knockdown *trp-2* channels in *Brugia*. Knockdown of *trp-2* by RNAi has previously demonstrated the key importance of TRP-2 in mediated diethylcarbamazine effects on motility as dsRNA *trp-2* animals showed reduced diethylcarbamazine sensitivity [4, 15]. We exposed dsRNA *trp-2* treated animals to 30 µM diethylcarbamazine and measured the Ca^2+^ signal. We observed that the Ca^2+^ was reduced but not inhibited in the *trp-2* dsRNA treated worms (Fig 5A; left panel, red trace & Fig 5B, red bar), with an average amplitude of 3% (3.4 %, ± 0.5 %, *n* = 3) compared to muscles not subjected to the dsRNA, 9 % (9.4 %, ± 1.6 %, *n* = 5: Fig 5A left panel, black line & Fig 5B brown bar). We observed no effect on the control CaCl_2_ signal in the dsRNA treated worms (26.8 %, ± 6.7 %, *n* = 3: Fig 5A right panel, red bar & Fig 5C, red bar) compared to our absent dsRNA samples (29.3 %, ± 8.8 %, *n* = 5: Fig 5A black line; right panel & C, grey bar). Using qPCR, we found that the dsRNA construct knocked down the level of *trp-2* by 83% (82.8 %, ± 2.3 %, *n* = 3) compared to 5.5% (5.5 %, ± 3.0%, *n* = 3) in LacZ controls (Fig. 5D). Our data suggests that TRP-2 is the major channel in mediating the diethylcarbamazine signal. Taken together our data suggests that the TRP-2 channel (an orthologue of vertebrate TRPC) is one of the major targets of diethylcarbamazine in adult *Brugia malayi* that contributes to the entry of Ca^2+^ into the muscle and mediates its effects on motility.

**Figure 5:**
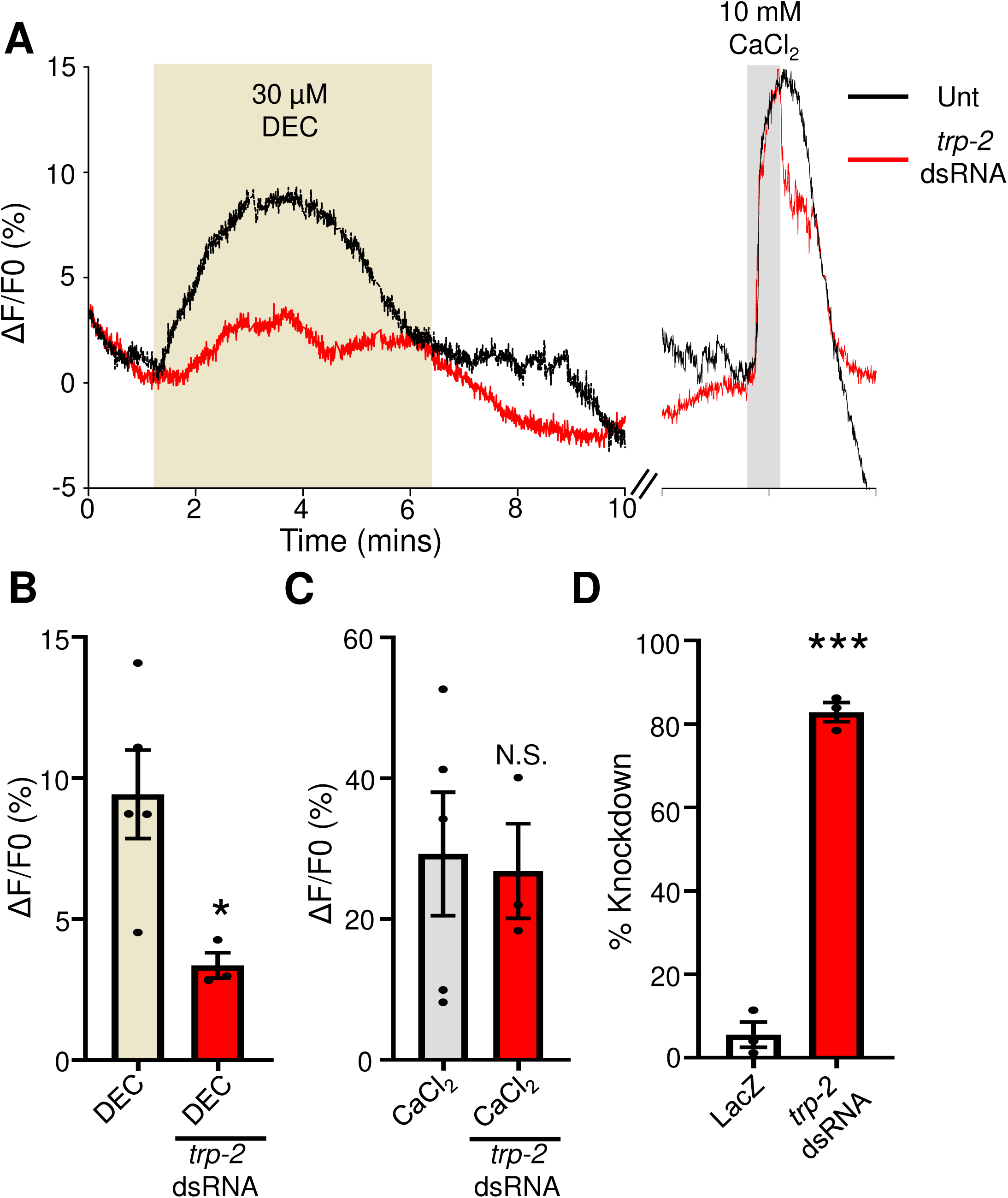
RNAi knockdown of *trp-2* reduces diethylcarbamazine responses. A) *Left panel:* Representative traces showing diethylcarbamazine induced Ca^2+^ responses in an untreated muscle (black line) and in a muscle treated with dsRNA for *trp-2* (red line). Light brown box highlights diethylcarbamazine application for 5 minutes. *Right panel:* Representative traces showing 10 mM CaCl_2_ induced Ca^2+^ responses in an untreated muscle (black line) and in a muscle treated with dsRNA for *trp-2* (red line). Grey box indicates 1 minute exposure to 10 mM CaCl_2_. B) Total amplitudes of Ca^2+^ fluorescence in response to 30 µM diethylcarbamazine in untreated muscles (light brown bar) dsRNA *trp-2* treated muscles (red bar). * Significantly different to untreated diethylcarbamazine. Diethylcarbamazine (DEC) vs dsRNA *trp-2* diethylcarbamazine, *P <* 0.029, *t* = 2.864, *df* = 6, unpaired *t*-test. C) Total amplitudes of Ca^2+^ fluorescence in response to 10 mM CaCl_2_ in untreated muscles (grey bar) and dsRNA *trp-2* treated muscles (red bar). N.S. Not significantly different to untreated muscle CaCl_2_. CaCl_2_ vs dsRNA *trp-2* CaCl_2,_ *P* < 0.855, *t* = 0.191, *df* = 6, unpaired *t*-test. *n* = 5 for diethylcarbamazine responses, and *n* = 3 for dsRNA *trp-2* diethylcarbamazine treatments and 10 mM CaCl_2_. D) qPCR analysis for *trp-2* expression in LacZ treated (control) and *trp-2* dsRNA treated *Brugia* represented as percentage knockdown. *** Significantly different to LacZ: dsRNA *trp-2* vs LacZ, *P <* 0.001, *t* = 20.260, *df* = 4, unpaired *t*-test. *n =* 3 for LacZ treated and *trp-2* dsRNA treated worms. All values represented as means ± SEM.

### Emodepside potentiates diethylcarbamazine Ca^2+^ signals

Emodepside is a current anthelmintic used in veterinary medicine that targets different parasitic nematodes and has significant potential for human use [26, 27]. Emodepside functions by targeting the Ca^2+^-activated K^+^ channel SLO-1 which causes paralysis of the nematode [11, 28]. We have demonstrated that emodepside produces flaccid paralysis in *Brugia,* by opening SLO-1 potassium channels and hyperpolarizes the muscle membrane of *Ascaris* [12, 13, 15]. Interestingly there is a synergistic relationship between emodepside and diethylcarbamazine resulting in potentiation of paralysis that is dependent on TRP-2 in *Brugia* [15] and which increases in the membrane potential hyperpolarization in the muscles of *Ascaris suum* [12].

We sought to determine if emodepside alone had any effect on the Ca^2+^ signal in our *Brugia* muscle cells. We predicted that there would be no effect of emodepside alone which by activating SLO-1 potassium channels would causes hyperpolarization [12] and thereby to close any voltage sensitive Ca^2+^ channels. We applied 1 µM emodepside and observed no changes in the Ca^2+^ signal amplitude (0.9 %, ± 0.4 %, *n* = 5) while there was no change in the control 10 mM CaCl_2_ Ca^2+^ signal (31.8 %, ± 8.1 %, *n* = 5: Fig 6A & B).

**Figure 6:**
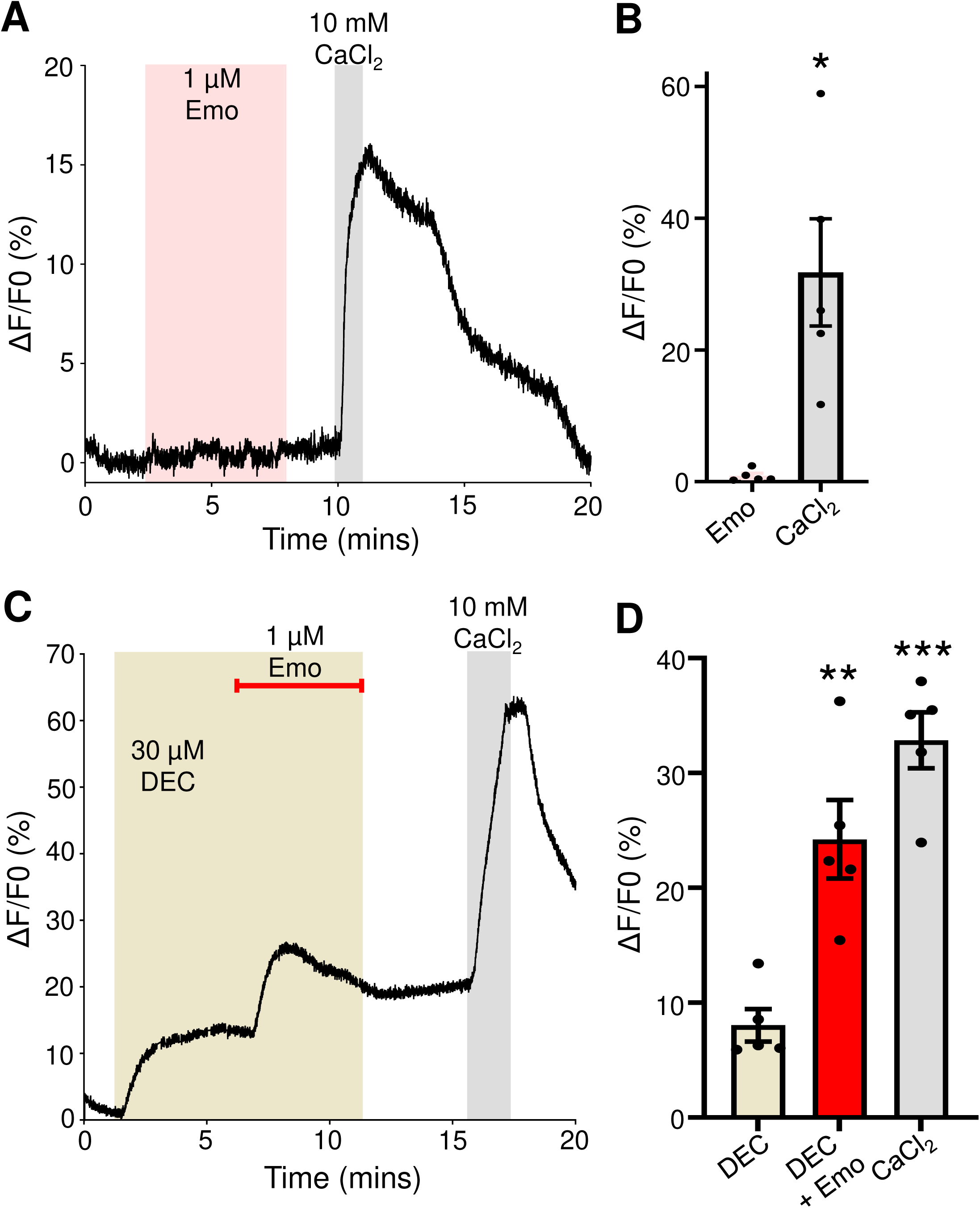
Combination of emodepside elevates diethylcarbamazine induced Ca^2+^. A) Representative trace in response to 5-minute exposure to 1 µM emodepside (pink box) followed by 1 minute exposure to 10 mM CaCl_2_ (grey box). B) Total amplitudes for emodepside (pink bar) and control 10 mM CaCl_2_ (grey bar) responses. * Significantly different to CaCl_2_. (Emodepside (Emo) vs CaCl_2_) *P <* 0.020, *t* = 3.745, *df* = 4, paired *t*-test. *n* = 5. C) Representative trace highlighting responses to 30 µM diethylcarbamazine (light brown box) for 5 minutes before the addition of 1 µM emodepside (red bar) with 30 µM diethylcarbamazine for 5 minutes. Samples were exposed to 10 mM CaCl_2_ (grey box) for 1 minute. D) Total amplitudes in response to 30 µM diethylcarbamazine (light brown bar), 30 µM diethylcarbamazine with the addition of 1 µM emodepside (red bar) and control 10 mM CaCl_2_ (grey bar). ** Significantly different to diethylcarbamazine. (Diethylcarbamazine (DEC) vs diethylcarbamazine (DEC) + emodepside (Emo)) *P <* 0.0066, *t* = 5.190, *df* = 4, paired *t*-test. (Diethylcarbamazine (DEC) vs CaCl_2_) *P <* 0.001, *t* = 10.72, *df* = 4, paired *t*-test. *n* = 5. All values represented as means ± SEM.

Subsequently, we applied 30 µM diethylcarbamazine for 5 minutes and observed the characteristic rise in Ca^2+^ that plateaued with an average amplitude of 8 % (8.0 % ± 1.4 %, *n* = 5: Fig 6C & D light brown bar) and then added 1 µM emodepside for 5 minutes on top of diethylcarbamazine: this generated a significant increase in the Ca^2+^ signal, 24% (24.2 % ± 3.4 % *n* = 5) which failed to decline after emodepside and diethylcarbamazine were removed (Fig 6C & D red bar). This observation may explain the persistence of reduced motility in parasites treated with both diethylcarbamazine and emodepside [15]. Our results illustrate a synergistic relationship between diethylcarbamazine and emodepside that together increases the entry of Ca^2+^ to disrupt the Ca^2+^ homeostasis within the muscles of *Brugia*.

## DISCUSSION

### Use of diethylcarbamazine and emodepside

Diethylcarbamazine, ivermectin and albendazole are recommended by WHO [1] for lymphatic filariasis mass drug administration (MDA) in areas without onchocerciasis. Diethylcarbamazine is a drug of choice for the treatment of lymphatic filariasis [7]. It kills microfilaria and is active against adult worms, although its effects are less pronounced. Diethylcarbamazine is generally well tolerated but side effects can occur when there are higher numbers of microfilaria in the blood. Diethylcarbamazine should not be used when humans are infected with onchocerciasis because it can make eye disease worse due to reactions of the larvae on the eye. There is also a concern of a possible severe reaction with individuals infected with *Loa loa* microfilaria and the development of encephalopathy.

Given that the existing registered antifilarial drugs (diethylcarbamazine, albendazole and ivermectin) do not kill all adult worms, there are concerns that those that survive administration of therapeutic doses will enhance the rate of development of resistance. Also, if a therapeutic mass drug administration (MDA) program were able to kill all adults so that microfilariae are no longer produced, elimination would be speeded up. This is because adults can survive for 6-8 years producing millions of microfilariae maintaining the life cycle [29, 30] if they are not eliminated.

Emodepside is an effective veterinary anthelmintic for intestinal nematode parasites that is undergoing Phase II clinical trials for onchocerciasis by DNDi that started in 2020. Emodepside opens nematode SLO-1 K^+^ channels [11, 13] and has inhibitory effects on the motility of different filariae (*Acanthocheilonema viteae, Brugia pahangi, Litomosoides sigmodontis, Onchocerca gutturosa* and *Onchocerca lienalis, Brugia malayi* and *Dirofilaria immitis*) when tested in preclinical investigations [31-33]. The effects of emodepside are dose-dependent, filarial species-specific-dependent and life-stage-dependent [34-39]. Adult *Brugia malayi* are a dose-limiting species because emodepside has the least potent effects on the adults of this filaria [31, 35]. Also, concentrations of emodepside may not be sufficient, for pharmacokinetic reasons, in regions where filarial parasites are located. The potential for diethylcarbamazine to potentiate the anthelmintic effects of emodepside [4,12,15] may prove useful in combination therapies.

### Interaction of diethylcarbamazine and emodepside involves TRP-2 and SLO-1 K channels

We have observed in *Ascaris suum* [12] and in adult *Brugia malayi* [4,15] that diethylcarbamazine potentiates the effects of emodepside. This potentiation was hypothesized to involve activation of TRP-2 channels that allow entry of Ca^2+^ to increase cytosolic Ca^2+^ and thereby increase emodepside activation of the Ca^2+^ - sensitive SLO-1 K channels. Here we have seen that application of diethylcarbamazine increases cytosolic Ca^2+^ concentrations by activation of the TRP-2 channel. The involvement of TRP-2 channels mediating the effects of diethylcarbamazine on cytosolic Ca^2+^ is revealed by inhibition of TRP-2 by the TRP-C antagonist, SKF96365 and by RNAi knockdown of *trp-2*. Furthermore, we were able to observe that not only does diethylcarbamazine increase cytosolic Ca^2+^ directly but that activation of SLO-1 K^+^ channels by emodepside enhances the Ca^2+^ response to diethylcarbamazine, Figs. 6 & 7. The opening by emodepside of potassium channels in the muscle membrane hyperpolarizes the muscle membrane and increases the driving potential for the entry of Ca^2+^ into the muscle cell through the TRP channels opened by diethylcarbamazine. Thus, there is a positive feed-back loop with diethylcarbamazine enhancing the effects of emodepside on SLO-1 K^+^ channels and emodepside enhancing the entry of Ca^2+^ produced by diethylcarbamazine Fig. 7. This positive feedback loop can explain the synergistic effect of diethylcarbamazine and emodepside on inhibition of motility [15].

**Figure 7:**
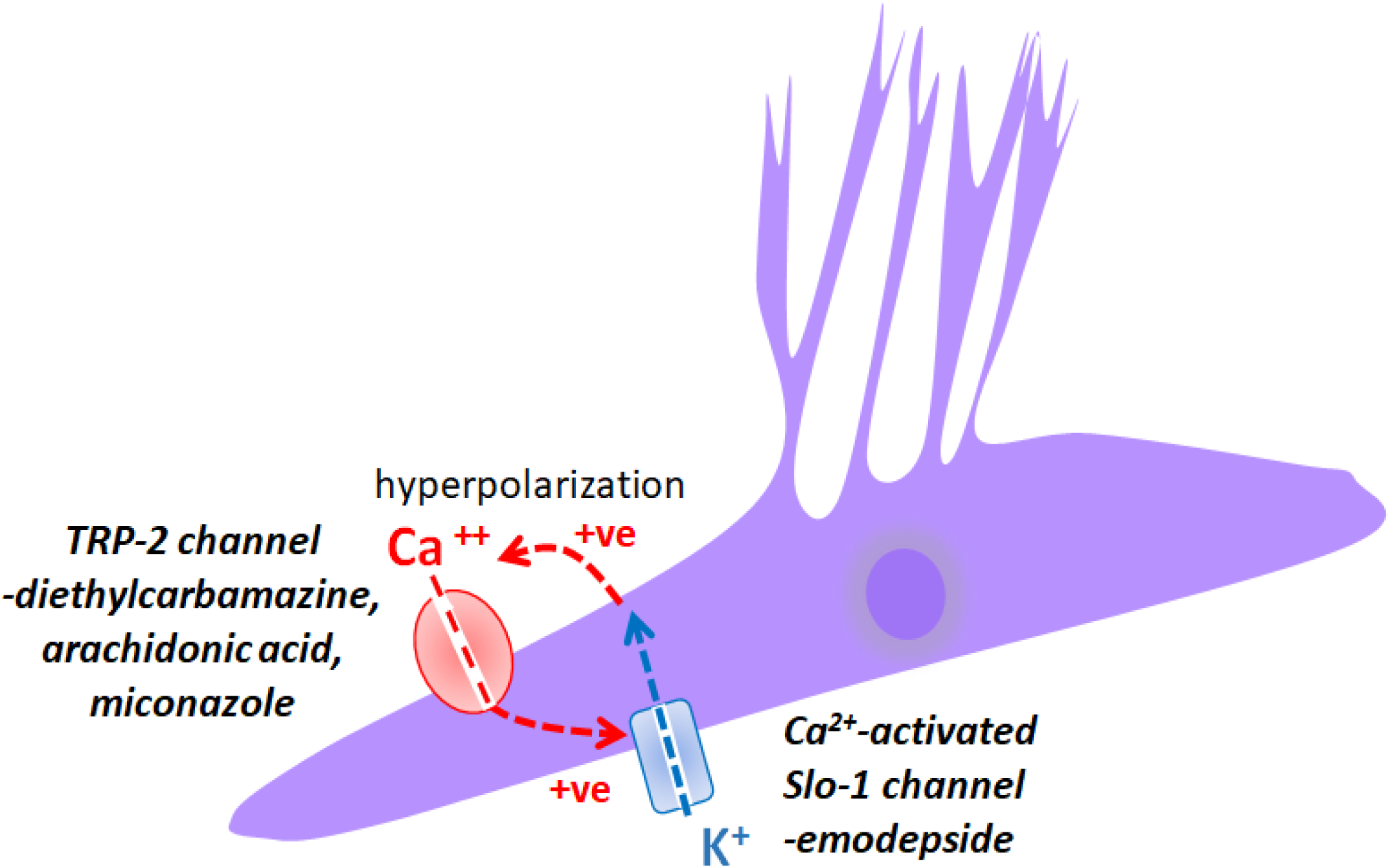
Summary diagram of the positive feed-back loop produced by diethylcarbamazine and emodepside. Diethylcarbamazine stimulates Ca^2+^ entry into *Brugia* muscle cells via TRP-2 and enhances the activation of SLO-1 K^+^ channels by emodepside that by hyperpolarizing the muscle cell further increases Ca^2+^entry. There is a synergistic relationship between the actions of diethylcarbamazine and emodepside produced by a positive feed-back loop. Diethylcarbamazine activates the TRP-2 channel and emodepside activates the SLO-1 K^+^ channels.

## Conclusion

We have seen that diethylcarbamazine has a rapid effect increasing cytosolic Ca^2+^ that is maintained during its application. The increased cytosolic Ca^2+^ will activate the contractile machinery of the muscle cells and limit the normal vibrating muscle activity that is seen in healthy worms. Emodepside enhances the effects of diethylcarbamazine with a positive feed-back loop: diethylcarbamazine opens TRP-2 channels, increasing cytosolic Ca^2+^ and activating SLO-1 K^+^ channels; and emodepside hyperpolarizes the membrane potential of the muscle, increasing the driving force for the entry of Ca^2+^. The combination of diethylcarbamazine and emodepside may be useful where the potency of either drug is limited by species of parasite, or location of the parasite.

## Acknowledgments

RJM is supported by NIH, the National Institute of Allergy and Infectious Diseases grants R01AI047194, R01AI155413 and the E. A. Benbrook Foundation for Pathology and Parasitology.

## Data Availability Statement

All relevant data are within the manuscript and its Supporting Information files.

---

**Supplementary Table 1:**
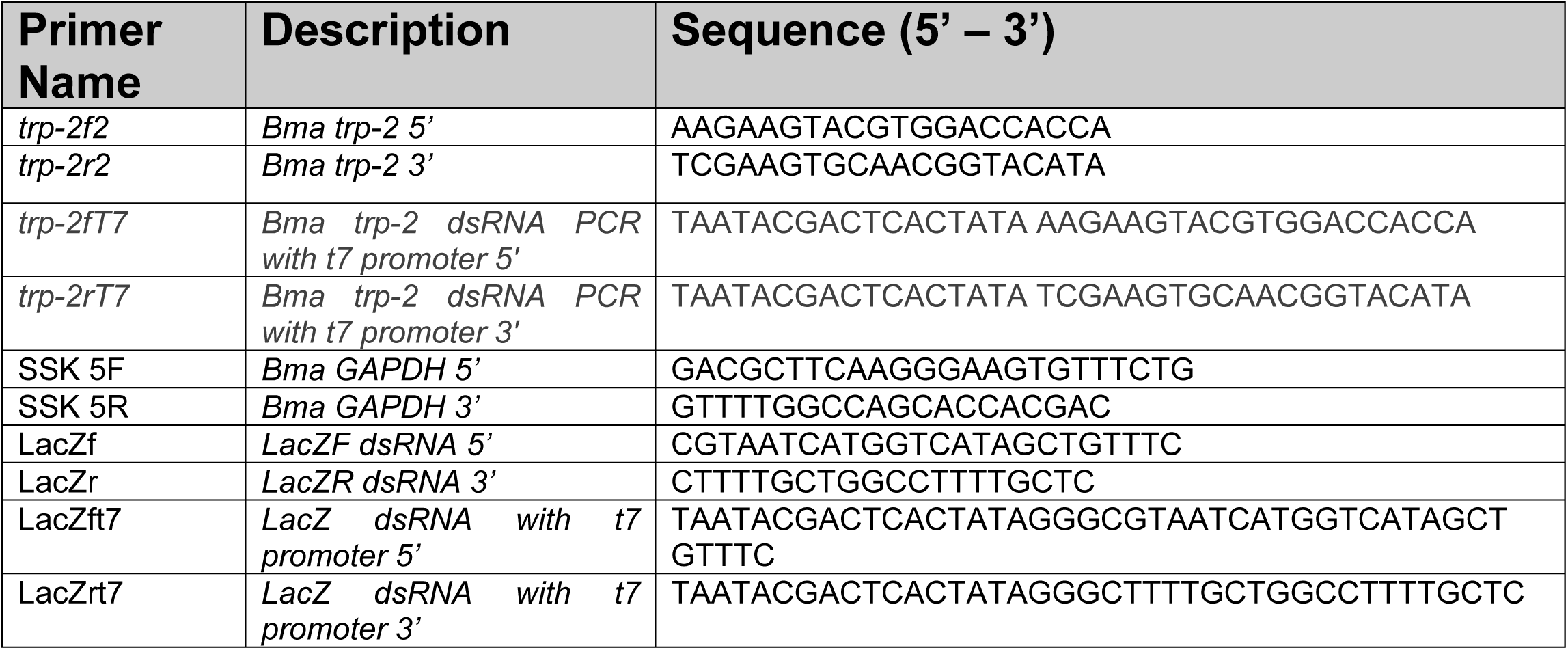
List of primers used in this study.

**Supplementary Table 2:**
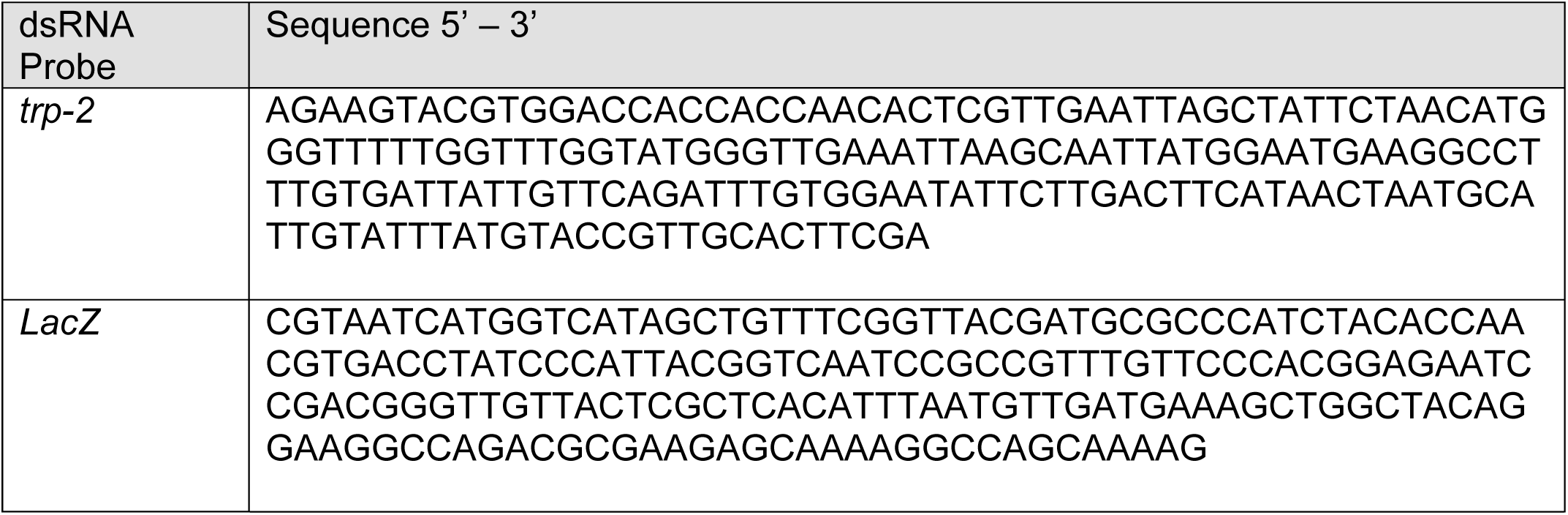
dsRNA probe sequences.

